# Evidence for strong fixation bias at 4-fold degenerate sites across genes in the great tit genome

**DOI:** 10.1101/436618

**Authors:** Toni I. Gossmann, Mathias Bockwoldt, Lilith Diringer, Friedrich Schwarz, Vic-Fabienne Schumann

**Author notes:** Correspondence: Toni I. Gossmann.

## Abstract

It is well established that GC content varies across the genome in many species and that GC biased gene conversion, one form of meiotic recombination, is likely to contribute to this heterogeneity. Bird genomes provide an extraordinary system to study the impact of GC biased gene conversion owed to their specific genomic features. They are characterised by a high karyotype conservation with substantial heterogeneity in chromosome sizes, with up to a dozen large macrochromosomes and many smaller microchromosomes common across all bird species. This heterogeneity in chromosome morphology is also reflected by other genomic features, such as smaller chromosomes being gene denser, more compact and more GC rich relative to their macrochromosomal counterparts - illustrating that the intensity of GC biased gene conversion varies across the genome. Here we study whether it is possible to infer heterogeneity in GC biased gene conversion rates across the genome using a recently published method that accounts for GC biased gene conversion when estimating branch lengths in a phylogenetic context. To infer the strength of GC biased gene conversion we contrast branch length estimates across the genome both taking and not taking non-stationary GC composition into account. Using simulations we show that this approach works well when GC fixation bias is strong and note that the number of substitutions along a branch is consistently overestimated when GC biased gene conversion is not accounted for. We use this predictable feature to infer the strength of GC dynamics across the great tit genome by applying our new test statistic to data at 4-fold degenerate sites from three bird species - great tit, zebra finch and chicken - three species that are among the best annotated bird genomes to date. We show that using a simple one-dimensional binning we fail to capture a signal of fixation bias as observed in our simulations. However, using a multidimensional binning strategy, we find evidence for heterogeneity in the strength of fixation bias, including AT fixation bias. This highlights the difficulties when combining sequence data across different regions in the genome.

## 1 INTRODUCTION

Estimating DNA sequence divergence between species is an important quantity in evolutionary analyses and population genetic approaches, such as for molecular dating, phylogeny reconstruction and the inference of selection. Several test statistics in population genetics that detect selection rely on an accurate reconstruction of the number of substitutions at putatively neutral sites, such as coding site (e.g. synonymous or 4-fold degenerate sites) or introns (Hudson et al., 1987; McDonald and Kreitman, 1991). For technical and computational reasons most popular models that estimate substitution rates assume that base composition is at equilibrium (Jukes and Cantor, 1969; Kimura, 1980; Felsenstein, 1981; Kimura, 1980, 1981). However, as there is a substantial base composition heterogeneity within and across genomes (Bernardi, 2000; Dreszer et al., 2007; Romiguier et al., 2010; Lartillot, 2013; Pouyet et al., 2017, 2018) this assumption is likely to be violated, and in fact base composition might change over time (Duret and Arndt, 2008).

Fundamental processes that contribute to heterogeneity in base composition over space and time are mutational bias and bias in the fixation probabilities of certain mutation types, such as due to selection, recombination, linkage or a combination of these factors (Eyre-Walker and Hurst, 2001). A particular example is the biased fixation probability that is caused by gene conversion of strong (G and C) over weak (A and T) base variants at heterozygous sites referred to as GC-biased gene conversion. It occurs during a repair induced gene conversion process that tends to preferably incorporate G/C nucleotides over A/T nucleotides during meiosis in many animal species (Duret and Galtier, 2009). Although the pronounced role of GC biased gene conversion as a major force is established, it is much less clear whether there is substantial variation in the extend of GC biased gene conversion across the genome and, if there is, how this is distributed across the genome. Several approaches have been developed that take heterogeneity in base composition into account when estimating substitution rates and branch lengths in phylogenies (Yang and Roberts, 1995; Galtier and Gouy, 1998; Dutheil and Boussau, 2008; Jayaswal et al., 2011) or by examining segregating variation (De Maio et al., 2013; Gleémin et al., 2015). However, it has been noted that accurately estimating sequence divergence can be difficult when GC content is not at equilibrium (Matsumoto et al., 2015).

Bird genomes are characterised by a high karyotype conservation across bird species pronounced by heterogeneity in chromosome sizes, with up to a dozen large macrochromosomes and many smaller microchromosomes (Ellegren, 2013). This heterogeneity in chromosome morphology is also in reflected in their genome composition features, with smaller chromosomes being gene denser, more compact and more GC rich relative to their macrochromosomal counterparts (Gossmann et al., 2014). As a consequence bird genomes provide an extraordinary system to study the evolution of GC hetereogeneity in a macrovolutionary context (Weber et al., 2014). To-date numerous bird genomes have been published (Zhang et al., 2014) with the genomes of chicken, zebra finch and great tit being among the best annotated high-quality bird reference genomes available (Hillier et al., 2004; Warren et al., 2010; Laine et al., 2016).

Here we infer heterogeneity in GC biased gene conversion rates across genes by estimating GC content evolution dynamics. We use a recently published method that accounts for nucleotide fixation bias when estimating branch length (Matsumoto et al., 2015). Using simulations we show that this approach works well and note that the number of substitutions along a branch are consistently overestimated when GC biased gene conversion is not accounted for in a stationary model. We use this predictable over-estimation as an indicator for the strength of GC dynamics across the genome and apply our new test statistic to data at 4-fold degenerate sites from three bird species - great tit, zebra finch and chicken. We use two binning strategies, current GC content of a focal species as applied previously and more complex clustering algorithm to estimate GC* (GC content at equilibrium). We find that binning according to current GC content, a frequently applied method (Bolivar et al., 2016; Corcoran et al., 2017), reveals little evidence for GC biased gene conversion across genes based on branch length estimations. In contrast, binning genes according to contemporary GC content of multiple species leads to a signal of GC and AT fixation bias as observed in our simulations, and suggests a substantially better model fit to the data. In conclusion there appears variation in the extent of strong fixation bias across genes with a signal for GC and AT fixation
bias.

## 2 MATERIALS AND METHODS

### 2.1 Sequencing data and phylogeny

We obtained sequencing data for coding genes from three bird species: great tit (Laine et al., 2016), zebra finch (Warren et al., 2010) and chicken (Hillier et al., 2004) - three of the best annotated and most studied bird genomes (Laine et al., 2018) currently available along with the high quality collared flycatcher genome (Ellegren et al., 2012; Kawakami et al., 2014) that was not considered here. An alignment pipeline was applied as described in Corcoran et al. (2017) from which we extracted aligned 4-fold degenerate sites only, as GC-biased gene conversion is supposed to act in particular on these sites (Bolivar et al., 2016). Since this is the only type of sites in this study and to improve readability, we refer to GC4 (GC content at 4-fold degenerate sites) as GC. We estimated branch lengths in a star like phylogeny based in a stationary model using baseml (Figure 1A) using the concatenated 4-fold sites alignments. These branch length estimates were then used to construct an approximated ultrametric tree as the underlying tree model for the nucleotide sequence simulations (Figure 1B).

**Figure 1.**
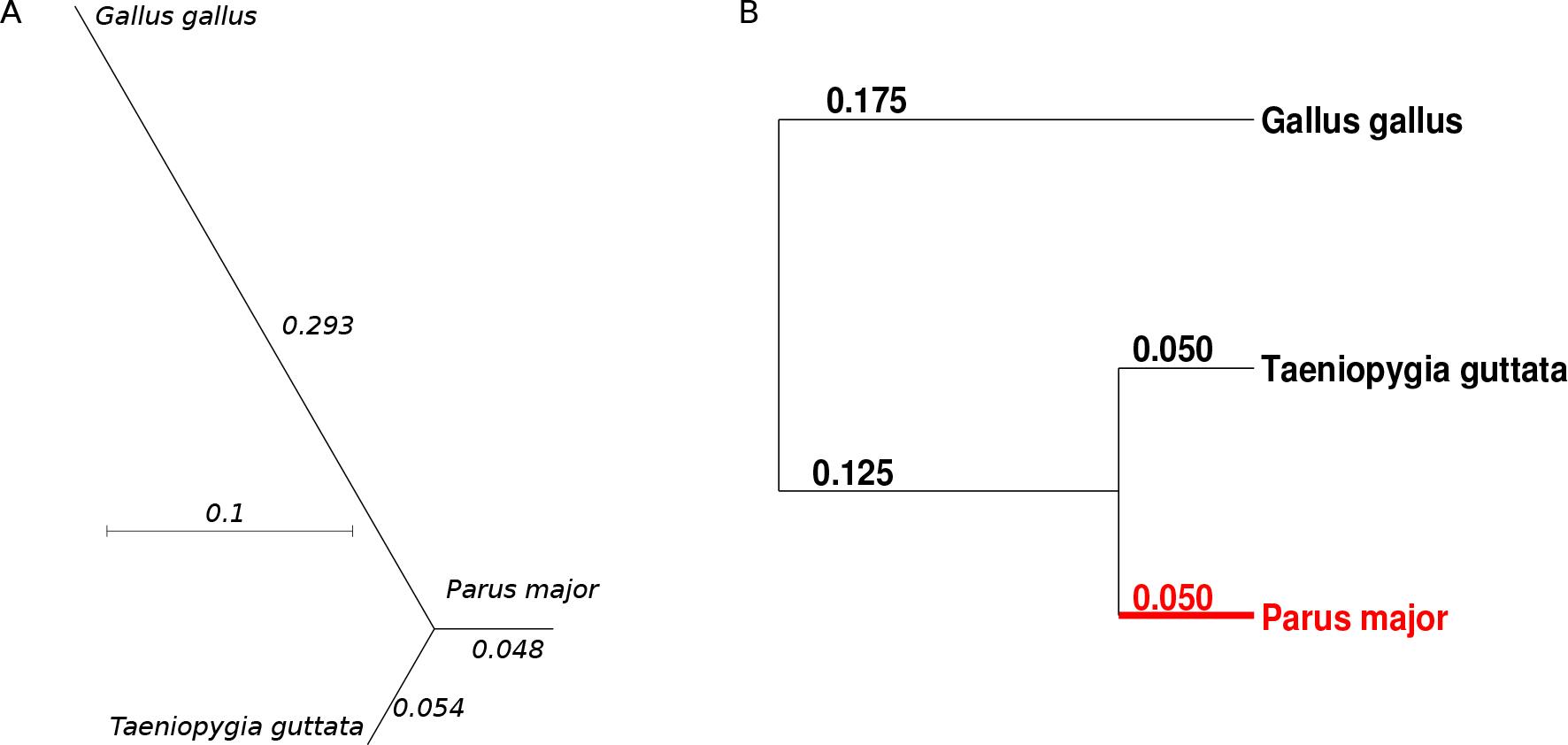
Three species (*Parus major, Taeniopygia guttata, Gallus gallus*) tree topology and (A) branch estimates from a stationary GTR model using baseml and (B) the ultrametric branch lengths assumed for the sequence simulations. The terminal branch in red was assumed to be evolving under non-stationary GC content.

### 2.2 Sequence simulation

We used INDELIBLE (Fletcher and Yang, 2009) to simulate nucleotide based sequence divergence with an underlying ultrametric tree topology as an estimate for branch length (Figure 1B). We assumed an HKY model (Hasegawa et al., 1985) with *κ* = 2.5 for the entire tree except for one of the shorter terminal branches for which we assumed a non-stationary model to simulate a non-stationary GC fixation bias. For this we used the UNREST substitution rate model in INDELIBLE (option 16) assuming following general matrix Q (Fletcher and Yang, 2009):

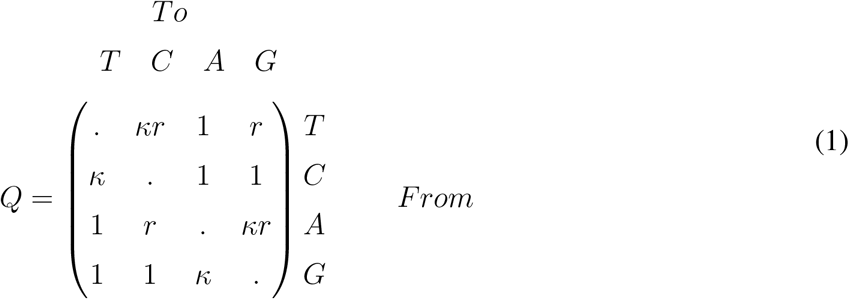

and define *r* = 1 + *B*/10 where B denotes the strength of GC fixation bias in the non-stationary process with an initial state frequency of *πA* = *πC* = *πG* = *πT* = 0.25. *B* values > 0 will result in a GC fixation bias while −10 < *B* < 0 values will result in an AT fixation bias. Please note that the commonly population size scaled gene conversion rate *B* = 4*N*_*e*_*b* (Nagylaki, 1983) is different from the B used here. We did not consider indels in the model.

### 2.3 Branch length estimation

Multiple processes can lead to fixation biases and their relative contributions are somewhat unknown. Here we assume that large scale variation in the fixation bias on 4-fold sites is largely driven by GC fixation bias which is not unrealistic (Smith et al., 2018). We repeated the forward simulations 100 times for each parameter set and estimated branch lengths in our tree by applying a method developed to reconstruct ancestral sequences when patterns of nucleotide substitutions are non-stationary Matsumoto et al. (2015) as implemented in PAML 4.8 (Yang, 2007) (i.e. model = 7, nhomo = 4, fix_kappa = 0). We also applied a simpler, homogeneous model (i.e. model = 7, nhomo = 1) using an unrooted tree topology that does not account for GC fixation bias. Log likelihoods were obtained from the model estimate and when obtained from binned data, summed across bins. The Akaike information criterion (AIC, Akaike (1974)) was used to assess model fit to compare different binning strategies and cluster numbers.

### 2.4 Binning strategy

We used two different binning strategies to combine data across genes. As contemporary GC content is relatively easy to measure we focused on current GC content per gene and clustered data using the k-means algorithm implemented in the scipy python package (kmeans2). We either used contemporary GC content of a single focal species which is comparable to equal binning sizes (Boĺivar et al., 2016; Corcoran et al., 2017) as well as multivariable clustering using contemporary GC content for each species. We note that other binning strategies may be applicable.

## 3 RESULTS

### 3.1 Sequence simulations

We conducted nucleotide forward simulations to generate non-stationary GC content in a terminal branch of an ultrametric three species tree using a customized substitution rate matrix with INDELIBLE. For simplicity reasons we assumed a phylogeny with ultrametric distances (Figure 1) although this is not a general restriction of the model. We simulated DNA stretches of 100 Kb without indels and applied two model tests in PAML to obtain branch length estimates. A simpler model that assumes stationary base composition (GTR, and an underlying unrooted tree) and a more complex model that incorporates non stationary base composition (GTR-NH, with a rooted tree). By that the more complex model should be able to capture GC or AT fixation biases in the terminal branch while the simpler model should not. Simulations were repeated 100 times and median estimates were obtained for varying strengths of GC fixation bias or sequence lengths.

#### 3.1.1 Inferring non-stationary GC composition

First we simulated DNA sequences under varying level of GC fixation bias (Figure 2). We confirm, as expected, that violating the assumption of stationarity in a phylogenetic model will lead to the parameters being estimated inaccurately. We observe an overestimation of the branch length with increasing GC fixation bias (Figure 2A) when the simple GTR model was used. If we estimate branch length in a model that accounts for fixation bias (GTR-NH) we can, however, accurately capture the correct branch length, even when GC fixation bias is extreme. We also note that there is an apparent discrepancy between the branch lengths estimated from the two models that is linear to the extent of fixation bias simulated (Figure 2A). Hence, the deviation in branch length estimates between the two models may be used as a proxy for the strength of fixation bias. To understand the base composition dynamics it is noteworthy that even with an enormous fixation bias (B = 59, the largest *B* value simulated here) GC content increases only moderately and GC content evolution is far from its equilibrium (Figure 2B). This is because of the relatively short branch length (however biological meaningful) considered here. This illustrates that the strength of fixation bias may be not correlated to the current GC content.

**Figure 2.**
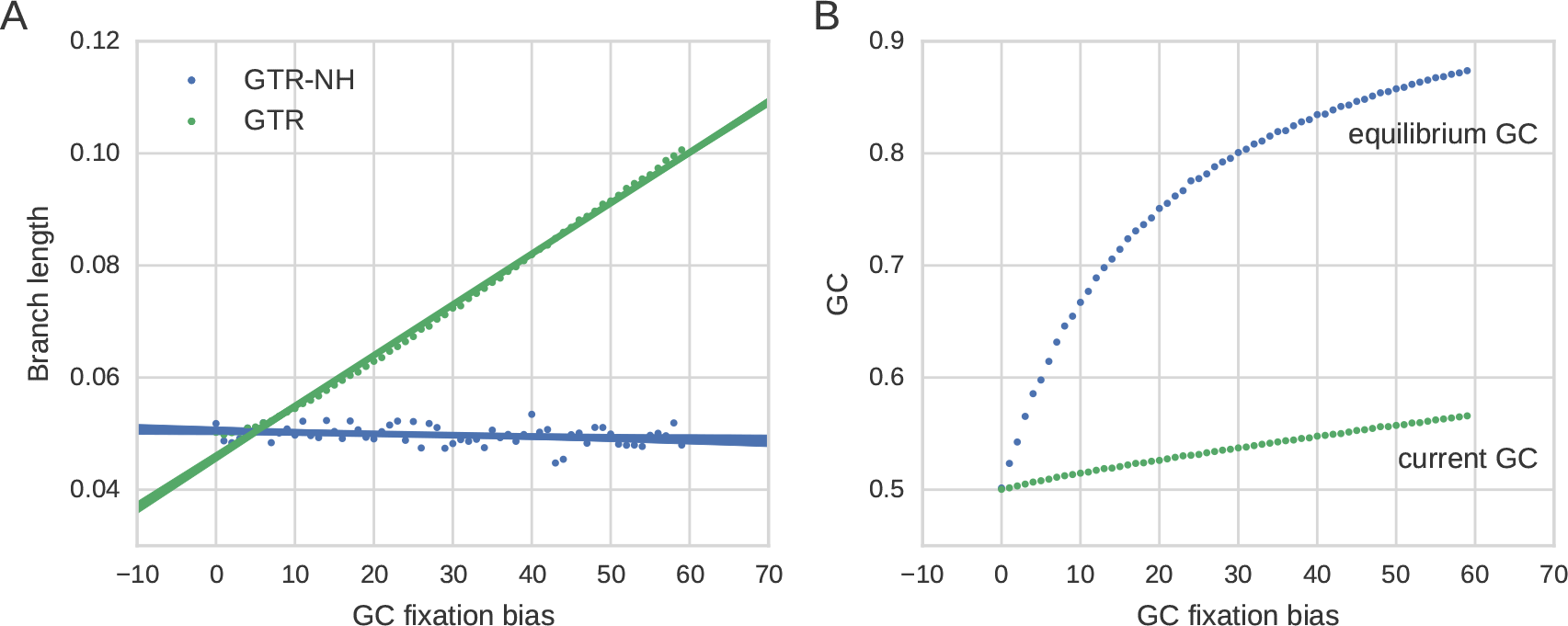
Features of GC content evolution in a terminal branch simulated under GC fixation bias. The terminal branch length under consideration is set to 0.05. (**A**) Estimated branch lengths under varying strength of GC fixation bias for stationary and non-stationary models. The solid line and associated confidence intervals denote the best fit of a linear regression model as implemented in the python function lmplot as part of the seaborn and matplotlib packages. (**B**) GC content evolution, shown is equilibrium GC (GC*) estimated from the GTR-NH model and GC content at the tip of the branch (current GC). Because of the short branch length GC content is far from its equilibrium.

#### 3.1.2 Parameter estimation and non-stationary GC composition in a non-focal branch

It was recently suggested that the effect of GC on branch length depends on the composition of the non-focal branches, as the stationary model estimates GC* from all branches (Guéguen and Duret, 2018). To study our model regarding the behaviour of non-focal branches we focused on the terminal sister branch (e.g. zebra finch lineage). We observe an overestimation of the branch length estimates in the GTR model with increasing GC fixation bias of the focal branch (Figure 3), although this effect appears to be modest and non linear. We also conducted additional simulation where we assumed the same extent of GC fixation bias for the focal branch and its sister branch (Figure 4) and find that parameter estimates are very similar to the case when GC composition is assumed to be stationary in the sister branch. Although we show only a modest effect of the non-focal branch in our simulation setup, this does not exclude the possibility of a more complex interplay between focal and non-focal branches when the underlying phylogeny is more complex as reported by Guéguen and Duret (2018).

**Figure 3.**
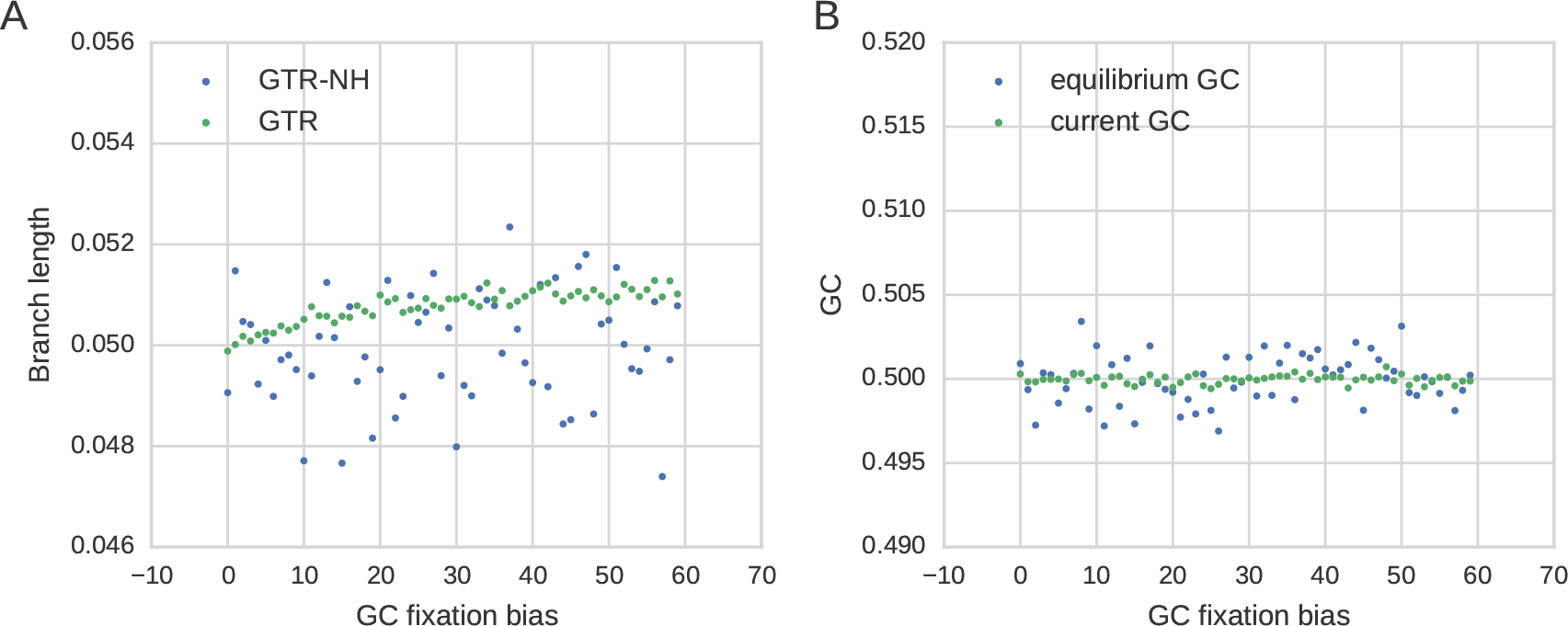
**Features of GC content evolution in the non-focal terminal branch** simulated under stationary GC when the focal branch is simulated with different levels of GC fixation bias. Terminal branch length under consideration is set to 0.05 in the simulations. (**A**) Estimated branch lengths under varying strength of GC fixation bias in the focal branch for stationary and non-stationary models. (**B**) GC content evolution, shown is equilibrium GC (GC*) and GC content at the tip of the branch (current GC).

**Figure 4.**
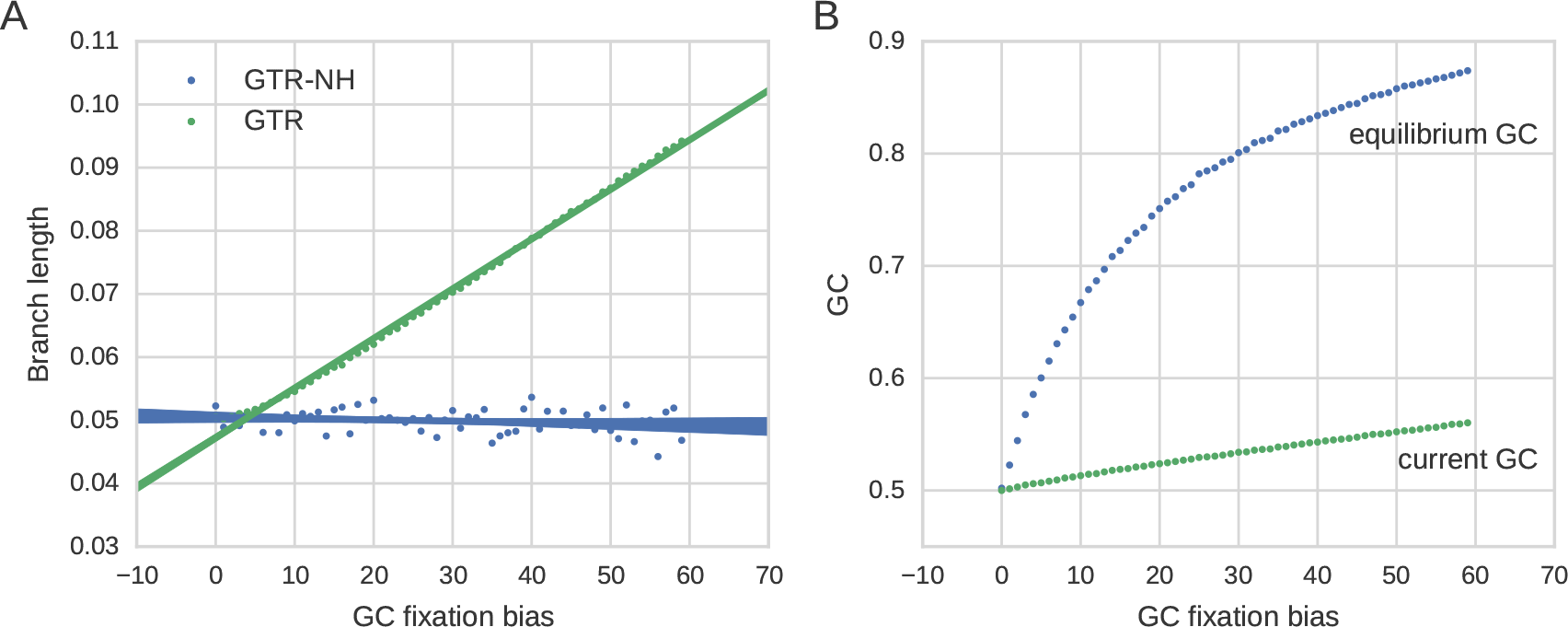
Features of GC content evolution in a terminal branch simulated under GC fixation bias when both terminal branches (e.g. great tit and zebra finch lineage) undergo the same level of GC fixation bias. The terminal branch length under consideration is set to 0.05. (**A**) Estimated branch lengths under varying strength of GC fixation bias for stationary and non-stationary models. The solid line and associated confidence intervals denote the best fit of a linear regression model as implemented in the python function lmplot as part of the seaborn and matplotlib packages. (**B**) GC content evolution, shown is equilibrium GC (GC*) and GC content at the tip of the branch (current GC). Because of the short branch length GC content is far from its equilibrium.

#### 3.1.3 Inferring non-stationary GC composition from limited data

We have shown that GC dynamics can be accurately captured when GC fixation bias is spatially homogeneous. To determine how much sequence information is necessary to accurately predict fixation bias, we conducted simulations of different sequence lengths with no, moderate and strong GC fixation bias (Figure 5). Under the assumption that the difference in branch length estimates between the GTR and GTR-NH model are a good proxy for determining the extend of GC sequencing bias, we find that fixation bias can be predicted very well. However, when sequence length is very short, the GTR-NH tends to mis-estimate the branch length and suggest that branch length can be accurately estimated when sequence are > 20*k*B.

**Figure 5.**
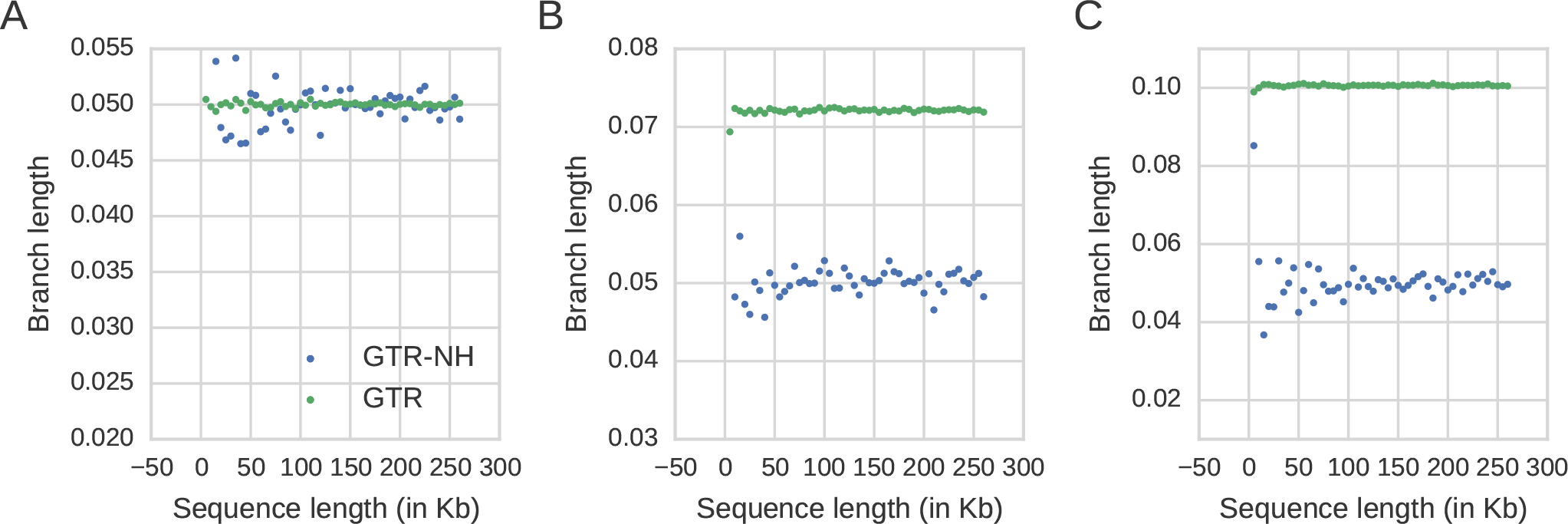
Comparison of a stationary (GTR) versus nonstationary model (GTR-NH) estimated from simulated data with varying sequence length. (A) When the transition matrix is symmetric (B = 0) (B) When GC fixation bias is moderate (B = 20) and (C) When fixation bias is strong (B = 59).

### 3.2 Application of non-stationary model to real data

As GC fixation bias is potentially correlated to GC content (Weber et al., 2014), sequence binning according to current GC content is a common method when gene sets of different strength of recombination are considered (Bolivar et al., 2016; Corcoran et al., 2017). Indicative of large scale GC composition dynamics at 4-fold degenerate sites stems from the per gene GC content distribution, which appears remarkably different between the chicken and passerine genomes (Figure 6). However, as the GC content distributions for the two passerine species appear very similar, the GC dynamics are potentially more subtle and difficult to infer. Here, to infer the GC content dynamics since the split of great tit lineage from the zebra finch lineage, we applied two kmeans clustering approaches to bin genes based on their contemporary GC content. First, we adopted the approach of Bolivar et al. (2016) of equal sized bins and applied a kmeans clustering on the GC content at the terminal branch (i.e. GC content of the great tit genes) per gene with varying cluster size. Second, we used a multidimensional kmeans algorithm (kmeans multidim) that takes the GC content per gene of all three species into account. The differences between these two approaches are illustrated in Figure 7 for an arbitrary cluster size of 20. In particular the cluster assignment for the chicken sequences differs between these two approaches.

**Figure 6.**
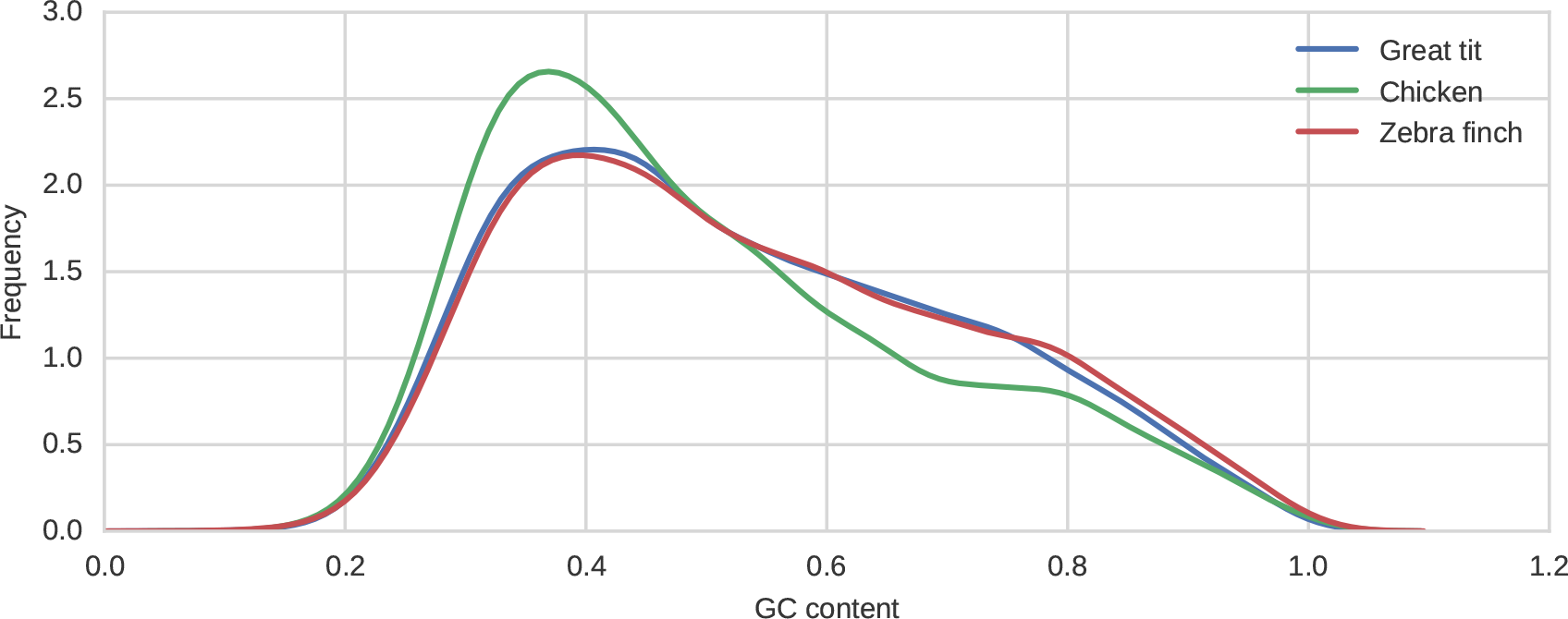
Distribution of GC4 content per gene for the three bird species under consideration. Shown are the kernel density estimations of the histograms as implemented in the python package seaborn.kdeplot.

**Figure 7.**
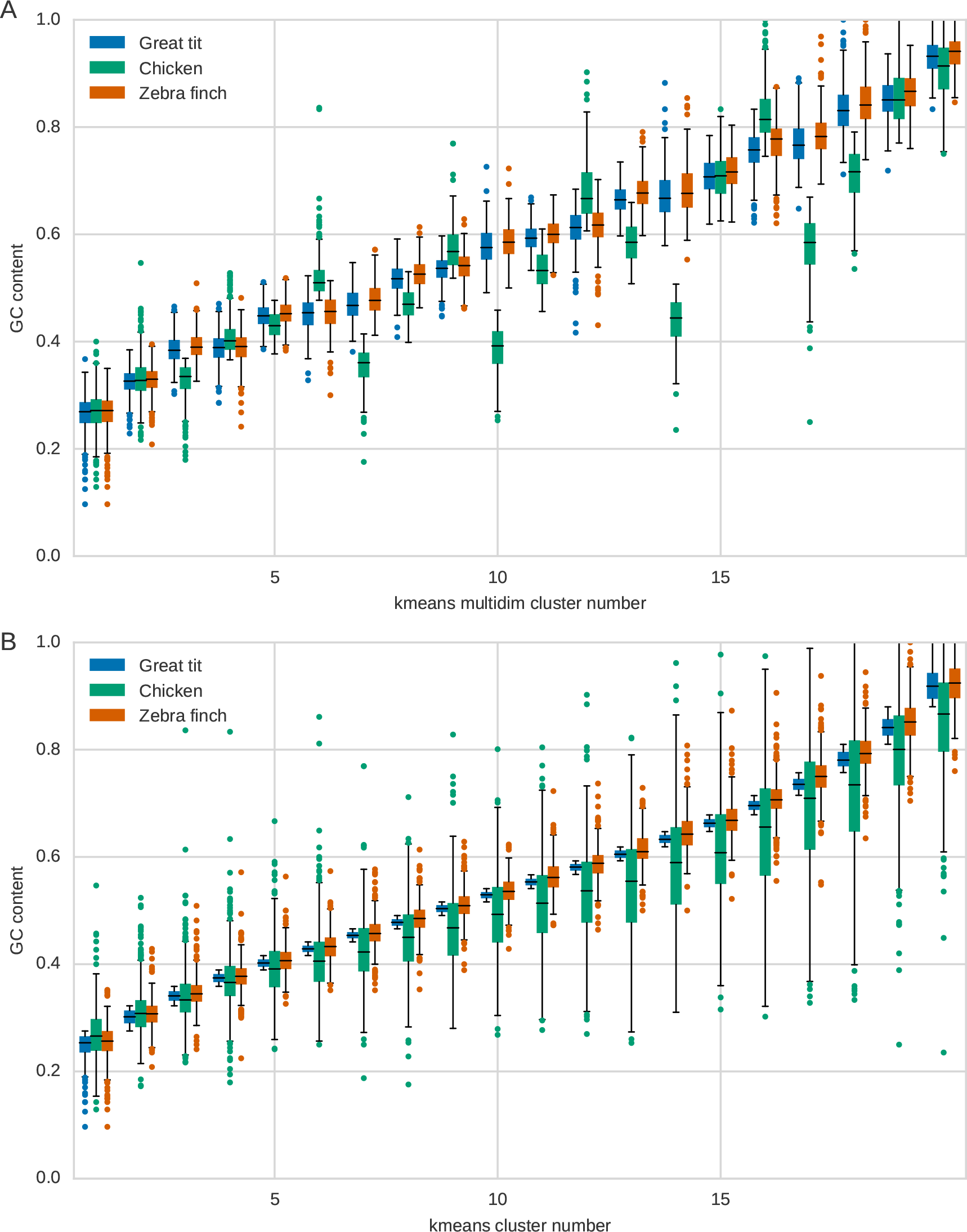
Boxplots of GC content per cluster for each species after applying a kmeans clustering approach using a pre-defined cluster number of 20 for (A) multidimensional kmeans clustering and (B) one-dimensional clustering.

We then applied the GTR-NH model to clustering outcomes and identified the best clustering outcome by comparing the AIC of the combined GTR-NH results. We find that the multidimensional clustering gives a much better fit to the data than clustering according to terminal GC content only (Figure 8A). We determine an optimal cluster size of 36 for the one-dimensional clustering and 187 for the multidimensional binning, but note that these numbers may vary because kmeans is implemented as a heuristic clustering approach. Results are qualitatively very similar across different cluster runs.

**Figure 8.**
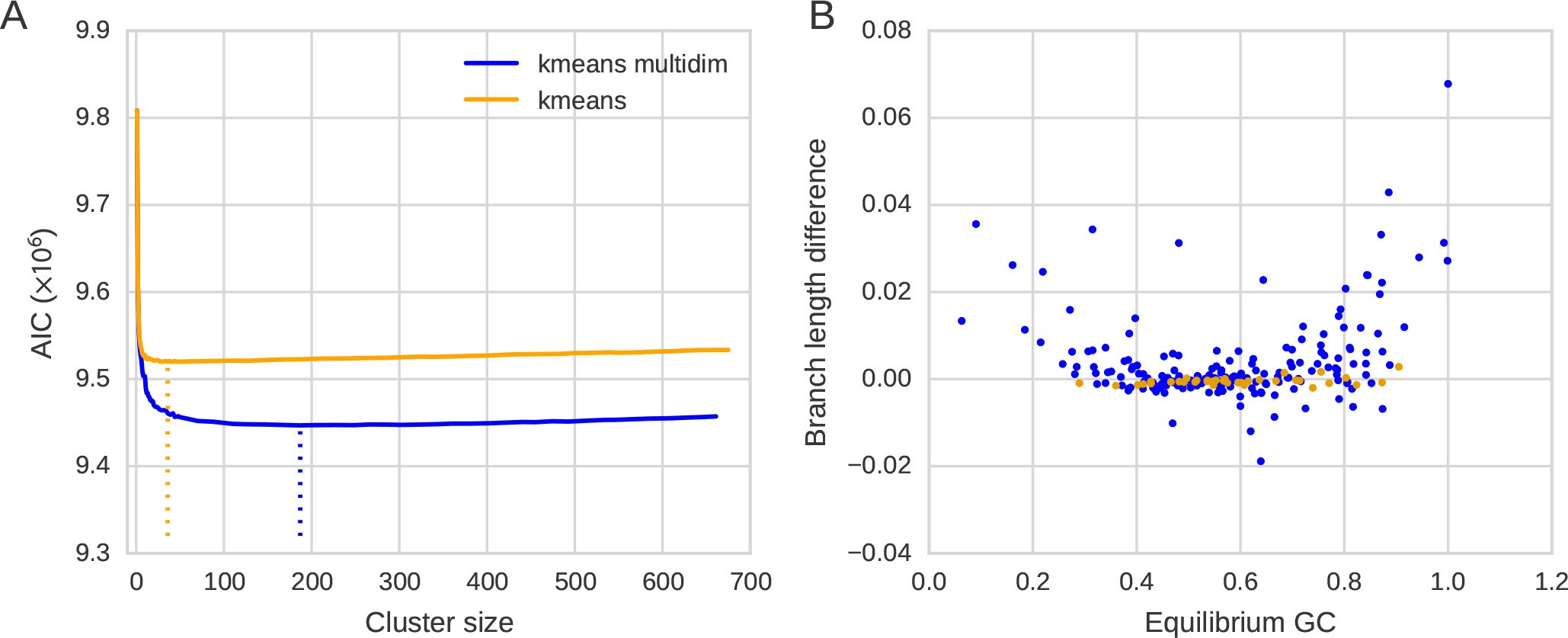
K-means clustering on GC content per gene. Clustering was conducted using a k-means implementation according to great tit GC content (kmeans) and according to terminal GC contents of the three species (kmeans multidim). (A) AIC of the GTR-NH model under varying cluster numbers. The optimal cluster numbers derived from the GTR-NH model fit are indicated with a dotted line (B) Branch length estimate difference and equilibrium GC for the optimal clusterings (kmeans and kmeans multidim).

To investigate how the two clustering strategies translate into capturing GC fixation bias we estimated branch lengths with the GTR and GTR-NH models to each cluster of the two optimal clusterings (kmeans and kmeans multidim). We then compared mean branch length differences between the GTR-NH and GTR models relative to the GC content at equilibrium obtained from the GTR-NH model (Figure 8B). For the one dimensional binning we observed very little discrepancy between branch length estimates (Figure 8B) at various levels of GC fixation bias. According to our simulations this may be observed when the extent of fixation bias is weak or when there is strong spatial heterogeneity in the extent of fixation bias. In contrary, for the multidimensional binning we see a discrepancy between branch length estimates for extreme GC and AT fixation biases, suggesting that both types of fixation biases occur in the genome, although more genes are prone to a GC fixation bias. We do not observe any functional enrichments of genes with either extreme GC fixation bias (GC*<0.2 and GC*>0.8, respectively) using a gene ontology enrichment analysis.

## 4 DISCUSSION

Here we have shown using simulations that taking non-stationary GC content into account when estimating branch lengths it is necessary and possible to capture the impact of nucleotide fixation bias. We also note that in our simulations fixation bias leads to a discrepancy in the branch length estimates between a stationary and non-stationary model, as previosuly reported (Matsumoto et al., 2015), and that this effect appears to be linear to the amount of fixation bias. We illustrate two major limitations of the non-stationary model applied here. First, it tends to be dependent on the GC dynamics at non-focal branches and secondly, it needs more data in comparison to a stationary model. Bearing these limitations in mind, we have applied the non-stationary model to 4-fold degenerate sites derived from gene alignments from great tit, zebra finch and chicken. Based on a Maximum-Likelihood approach we find that fixation bias can be potentially accounted for when subdividing the dataset into smaller bins. This yields better model fits according to AIC and a few bins are already sufficient to improve the fits substantially. This rough binning might suggest that there is large scale variation in the extent of GC fixation bias, but here we argue that it could also be simply driven by variation in the base composition at the ancestral or terminal node across loci.

To investigate whether there is truly variation in the fixation bias, we apply two different binning strategies to estimate fixation bias separately for smaller sets of genes which allows to include information on very short genes. We show that to accurately capture the role of fixation bias the method of clustering is crucial. A simpler one-dimensional binning according to current GC content for the terminal branch under consideration leads to a relatively low cluster number (i.e. 36 clusters). Moreover, for the estimated bins we fail to capture a signal of fixation bias based on branch length estimates that we observe in our simulations. This is also observed for larger cluster numbers using this clustering strategy (results not shown). Under such a simple clustering method, we find an almost perfect correlation between current GC content and estimated equilibrium GC content (Kendall *τ* = 0.99, P<<0.05) as observed by others (Weber et al., 2014). In the light of our simulations this suggests that it is difficult to capture true GC fixation bias across the genome when taking contemporary GC content of a single species as the only clustering variable into account.

We observe variation in the equilibrium GC content and branch length estimates when we use a multidimensional binning. In concordance with our simulations we observe an increased difference in the branch length estimates between GTR and GTR-NH model. Secondly, also with moderate equilibrium GC content (0.2<GC*<0.8) we observe differences in the branch length estimates between the two models. Our simulations suggests that little sequence data may lead to GTR-NH model to underestimate the true branch length. To check whether bins with little sequence data are driving the observed pattern, we correlated total sequence length per bin with current and equilibrium GC content as well as differences in branch lengths estimates. We do not find any significant correlation of sequence length and GC content and equilibrium GC content (Kendall *τ* = −0.0311, P = 0.53 and *τ* = −0.025, P = 0.6, respectively), but do find a significant correlation between sequence length and branch length estimate differences between the two models (Kendall *τ* = −0.26, P = 1.3 × 10^−7^). If we remove the top 20% of bins with extreme branch length difference, this relationship remains significant (Kendall *τ* = −0.22, P = 4.1 × 10^−5^) - suggesting that the pattern is not driven by extreme outliers. We also observe a significant correlation between contemporary GC content and equilibrium GC (Kendall *τ* = 0.48, P<<0.05), which is less pronounced than in the one-dimensional binning. It is however possible that our method misses fine scale variation in GC fixation bias, which we are unable to address in the model used due to lack of data. We also do not consider within gene variation in GC fixation bias as shown for plants (Glémin et al., 2014), although this aspect could be captured by modification to the binning strategy.

We have considered a model of spatial heterogeneity in GC dynamics, however, we have not taken into account temporal variation in the GC dynamics across the genome. Temporal GC dynamics are probably less likely to occur in comparison to mammalian genomes as birds lack the recombination hotspot protein PRDM9 (Singhal et al., 2015). However, unlike interchromosomal rearrangements intrachromosomal rearrangements are not uncommon in bird genomes (Romanov et al., 2014), suggesting that sudden changes in the recombination environment and hence the rate of fixation biases are possible. Evidence supporting this notion stems from the observation in our analysis that contemporary GC content is much less correlated with fixation bias when binning data with a multidimensional kmeans approach. It is also unclear whether we underestimate the amount of extreme GC bias, as we might miss genes of extreme GC composition in our dataset - a relatively large number of genes are not annotated in many bird genomes (Lovell et al., 2014; Hron et al., 2015; Botero-Castro et al., 2017), and this is likely to be an artefact of technical difficulties to sequence genomic reads with extreme nucleotide composition.

In conclusion, we have shown using simulations and real data analysis that care has to be taken when estimating branch length under the impact of fixation bias. As noted previously tends GC fixation bias lead to an overestimation of the true rate of fixation. We note that under moderate fixation bias this effect is relatively small. The suggested binning strategy may be useful when applying tests of non-neutral evolution across the genome.

## AUTHOR CONTRIBUTIONS

The author contributions are as follows. TIG designed the study, conducted the simulations and wrote the manuscript. LD contributed to the initial draft of the manuscript. LD, FS and VFS conducted simulations and contributed code.

## ACKNOWLEDGMENTS

We thank Henry Barton for commenting on an earlier version of this manuscript.

## DATA AVAILABILITY STATEMENT

An example script file for INDELIBLE used for this study can be found in the git hub repository https://github.com/tonig-evo/GCdym

